# Probing the ionotropic activity of the orphan glutamate delta 2 receptor with genetically-engineered photopharmacology

**DOI:** 10.1101/2020.05.14.093419

**Authors:** Damien Lemoine, Sarah Mondoloni, Jérôme Tange, Bertrand Lambolez, Philippe Faure, Antoine Taly, Ludovic Tricoire, Alexandre Mourot

## Abstract

Glutamate delta (GluD) receptors belong to the ionotropic glutamate receptor family, yet whether they actually form functional and physiologically-relevant ion channels in neurons remains a debated question. Here we used a chemo-genetic approach to engineer specific and photo-reversible pharmacology in the orphan GluD2 receptor. We incorporated a cysteine mutation in the cavity located above the putative ion channel pore, for site-specific conjugation with a photoswitchable ligand. We first showed that, in the constitutively-open GluD2 Lurcher mutant, current could be rapidly and reversibly decreased with light. We then transposed the cysteine mutation to the native receptor, to demonstrate with absolute pharmacological specificity that metabotropic glutamate receptor signaling opens the GluD2 ion channel in heterologous expression system. Our results assess the functional relevance of GluD2 ion channel and introduce an optogenetic tool that will provide a novel and powerful means for probing GluD2 ionotropic contribution to neuronal physiology.

---

Glutamate delta (GluD1 and GluD2) receptors are considered orphan because, while having a strong sequence homology with the other ionotropic glutamate receptors (AMPA, NMDA and Kainate), they are not activated by glutamate^1,2^. GluD receptors are both widely expressed throughout the brain, GluD1 predominating in the forebrain, and GluD2 being highly enriched in cerebellar Purkinje neurons^3,4^. Both GluD1 and GluD2 play a role in the formation, stabilization, function and plasticity of synapses^4-7^. Likewise, deletion of GluD1 or GluD2 genes in mouse results in marked behavioral alterations^8,9^, and mutations in human GluD1 and GluD2 genes have been associated with neurodevelopmental and psychiatric diseases^10,11^, attesting to their functional importance in brain circuits. Nevertheless, due to the absence of pharmacology, a detailed understanding of how GluD1/2 regulate specific neural circuits, and notably whether their ionotropic activity is involved, is lacking.

Although GluD1 and GluD2 exhibit a domain similar to the ligand binding domain (LBD) of other iGluRs^12^, no ligand has been found that directly triggers the opening of the pore. Yet, several observations indicate that the ion channel of GluD receptors may be functional. First, crystallization studies show that the LBD of GluD2 binds D-serine and glycine, and that these ligands induce “agonist-like” structural rearrangements in the LBD, even though they fail to evoke currents at wild-type (WT) GluD receptors expressed in heterologous expression systems^13^. Second, a point mutation (A654T) in GluD2 that causes the degeneration of cerebellar Purkinje neurons in *Lurcher* (Lc) mice confers constitutive ion flow^14,15^. Current through GluD2^Lc^ receptors is inhibited by pentamidine and 1-Naphthyl acetyl spermine (NASPM)^16,17^, pore blockers of NMDA and AMPA receptors, respectively. Furthermore, D-serine and glycine reduce the spontaneous currents of GluD2^Lc^, suggesting a coupling between the LBD and the channel^13,18^. Third, receptor chimeras containing the LBD of AMPA receptors and the membrane domain of GluD receptors show glutamate-induced currents^19^. Fourth, the GluD1/2 receptor channel can be opened following activation of type I metabotropic glutamate receptors (mGlu1/5), and these currents are almost completely blocked by NASPM and reduced by D-serine^20-22^. Finally, the slow excitatory postsynaptic currents observed in midbrain dopaminergic, dorsal raphe, and cerebellar Purkinje neurons, are abolished upon gene inactivation or expression of dominant-negative pore mutants of GluD1/2^20,22,23^. All these findings indicate that GluD receptors likely possess a functional ion channel pore. Yet, a direct evidence for ionotropic activity of GluD in neuronal setting is lacking, due to the inability to specifically and acutely block GluD conductance.

To fill this gap, we bestowed light-sensitivity to the GluD ion channel pore using an optogenetic pharmacology approach^24^. We incorporated a cysteine point mutation at the surface of GluD2, right above the hypothetical channel lumen, onto which can be anchored a photoswitchable tethered ligand (PTL). Light is then used to modify the geometry of the PTL, thereby presenting/removing the ligand to/from the channel, resulting in optical control of ionotropic activity. Here we demonstrate rapid and reversible, optical control of ion current through a cysteine-substituted GluD2 receptor. This novel tool, called light-controllable GluD2 (LiGluD2), allows rapid, reversible and pharmacologically-specific control of GluD2, and may help provide a mechanistic understanding of how this receptor contributes to brain circuits and behaviors.

Our approach to probing the functionality of the ion channel in GluD is to install a photo-isomerizable pore blocker at the extracellular entrance to the channel lumen (Figure 1A). The tethered ligand is site-specifically attached to a cysteine-substituted residue. In darkness or under green light, the PTL adopts an elongated shape and reaches the lumen, resulting in ion channel blockade, while under violet light, it switches to a twisted, shorter configuration, relieving blockade. Our design of the PTL was based on the chemical structure of pentamidine (Figure 1B), a pore blocker that efficiently blocks current through GluD2^Lc^ receptors^17^. The PTL, called MAGu, contains a thiol-reactive maleimide (M) moiety, a central photo-isomerizable azobenzene (A) chromophore, and a guanidinium (Gu) head group that resembles the amidinium groups of pentamidine (Figure 1C). MAGu was selected notably because its synthesis route has been described (referred to as PAG1c in the original article^25^). In aqueous solution, MAGu could be converted to its *cis* form using 380 nm light, and converted back to *trans* either slowly in darkness (t1/2 ∼ 20 min) or rapidly upon illumination with 525 nm light (Supp. Fig.1A-B), in agreement with previous reports^26^. To find the best attachment site for MAGu on GluD, we developed a homology model of the GluD2 receptor, based on the structure of the recently crystallized GluA2 receptor^27^ (see methods). Using this model, we selected a series of 15 residues, located on the peptide that links the LBD to the third transmembrane domain (TM3) that lines the channel lumen, for mutation to cysteine (Figure 1D-E).

**Figure 1:**
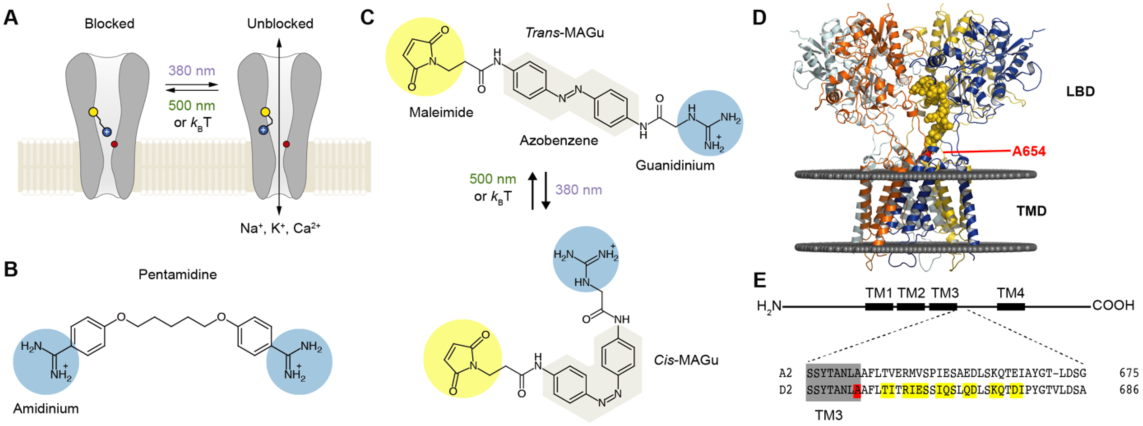
Optogenetic pharmacology strategy to probe the ionotropic activity of GluD receptors. A. GluD2 is genetically-modified to incorporate a cysteine residue (yellow) at the entrance to the pore, which serves as a handle for the covalent attachment of a synthetic, photoswitchable tethered ligand (PTL). Under green light (500 nm), the PTL adopts an elongated state and places its cationic head group in the lumen, resulting in ion channel blockade. Under violet light (380 nm), the PTL switches to a twisted, shorter form and unblocks the channel. The position of the Lurscher (Lc) mutation, which produces a permanently open channel, is depicted in red. B. Chemical structure of pentamidine, a non-selective iGluR blocker with two amidinium head groups. C. Chemical structures of the PTL MAGu in its *trans* (top) and *cis* (bottom) configurations. MAGu is composed of a cysteine-reactive maleimide group, a central azobenzene chromophore, and a guanidinium cationic head group. D. Molecular model of GluD2, based on the structure of activated GluA2 (5weo). Residues mutated to cysteine are depicted in yellow, while the Lc mutant is shown in red. E. Top, schematic representation of one GluD subunit, with its four transmembrane segments (TM). Bottom, sequence alignment between the mouse GluA2 and GluD2 receptors around the engineered mutations. TM3 is shown in grey, the 15 residues mutated to cysteine in yellow, and the position of the A654T Lc mutation in red.

Since no known ligand directly gates the ion channel of GluD2, we used a Lc mutant, L654T, which displays a constitutively open channel^14,15^, for screening the 15 single-cysteine mutations. Accordingly, we found that heterologous expression of GluD2-L654T, but not of the wild-type (WT) protein, in HEK cells produces large currents that reverse at membrane potential close to 0 mV and are reduced by externally-applied pentamidine (Figure 2A). Subtracted Lc current showed clear rectification at positive potentials, as reported with the blockade by NASP another GluD blocker^16^. Therefore, the L654T Lc mutant was subsequently used as a screening platform to find the best attachment site for MAGu on GluD2. Each of the 15 residues identified in Fig. 1D were mutated individually to cysteine on the L654T background, and tested using patch-clamp electrophysiology. Cells were treated with MAGu (20 µM, 20 min) and Lc currents were measured in voltage-clamp mode (−60 mV) under different illumination conditions to toggle MAGu between its *cis* and *trans* states. Current through L654T was not affected by light, indicating that MAGu has no effect on this Lc channel. In contrast, we found several cysteine mutants for which current was significantly larger under 380 than under 535 nm light, and one mutant (Q669C) for which there was a tendency for “reverse photoswitching”, i.e. larger currents under 535 than under 380 nm light (Figure 2B). We then quantified the degree of photoswitching by comparing the block in darkness (*trans* state) to the block evoked by pentamidine. We excluded from the analysis mutants that displayed no pentamidine-decreased leak current (i.e. mutants for which pentamidine block was significantly smaller than that observed on A654T, Fig. Sup 2A-B), because they were likely either not expressed or not functional. Photoswitching was significant for Q666C, Q669C, D670C, Q674C and I677C, suggesting that MAGu covalently reacted with these cysteine mutants and that, once tethered, it could modulate current in one of its conformer (Figure 2C). Importantly, photomodulation was absent in the control A654T and the other cysteine mutants, indicating that the effect of light is specific to the attachment of MAGu to the above-mentioned cysteine mutants. From a structural point of view, the photocontrollable mutants are all located at the very top of the linker, i.e. further away from the TMD compared to other tested residues (Figure 2D).

**Figure 2:**
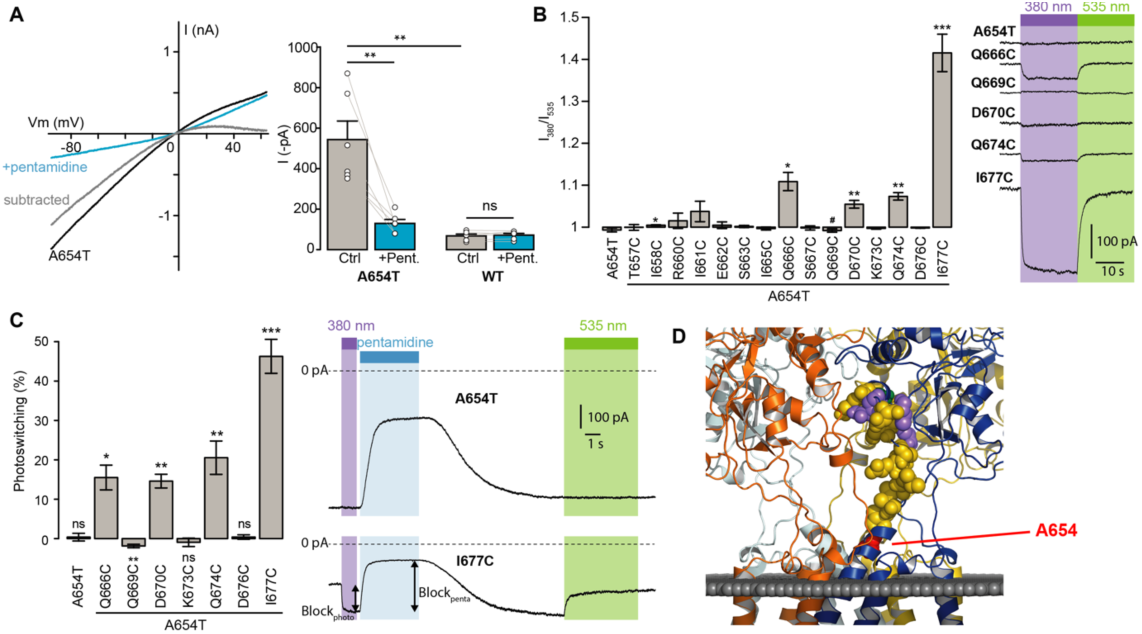
Screening of the fifteen single-cysteine mutants engineered on the Lc background. A. Left, representative current-voltage relationship for GluD2-A654T Lc, with (blue) and without (black) pentamidine (100 μM). The subtracted current (grey) shows clear inward rectification and block at positive voltages. Right, currents recorded at −60 mV were larger for GluD2-A654T than for WT (n=6 cells, p = 0.0033, two-sample t-test), and were strongly reduced with 100 μM pentamidine for A654T (n=6 cells, p = 0.0045, paired t-test) but not for the WT GluD2 (n = 6 cells, p = 0.34, paired t-test). B. Left, ratio of the currents recorded at −60 mV under 380 and 535 nm light, for A654T and the fifteen cysteine mutants engineered on the A654T background (n = 3-8 cells, one-sample t-test, or Wilcoxon when normality is not verified, compared with a theoretical mean value of 1). Right, representative change in holding current when switching between dark, 380 and 535 nm light, for A654T and for 5 mutants that show modulation of the holding current when switching between 380 and 535 nm light (p< 0.05 except for Q669C where p = 0.09). C. Left, percent photoswitching (PS = Block_photo_/Block_penta_) for A654T and the cysteine mutants that show both clear leak current and pentamidine block. Photoswitching reached 15.5 ± 3.1 % for Q666C (p=0.016, n = 4), −1.9 ± 0.4 % for Q669C (p=0.008, n =6), 14.6 ± 1.7 % for D670C (p=0.003, n=4), 20.5 ± 4.2 % for Q674C (p=0.008, n=5) and 46.3 ± 4.3 % for I677C (p = 1.34E-05, n= 8, one-sample t-test, or Wilcoxon when normality is not verified, compared with a theoretical mean value of 0). Photoswitching was absent in the control A654T mutant (0.37 ± 0.9 %, p = 0.72, n = 5) and in the other two cysteine mutants K673C (−0.9 ± 1.06 %, p = 0.43) and D676C (0.39 ± 0.57 %, p = 0.53). Right, representative current (Vm = −60 mV) recorded for A654T and A654T-I677C in darkness, under 380 and 535 nm light, with and without pentamidine (100 μM). D) Model of GluD2 showing the location of A654 (red), Q666, D670, Q674 and I677C (violet), Q669 (green), and all the non-photocontrollable cysteine mutants (yellow). Data are presented as mean value ± sem.

We then selected the best mutant GluD2-L654T-I677C for further characterization. Photoregulation was fully reversible over many cycles of 380 and 535 nm light (Fig. 3A), in agreement with the fact that azobenzenes photobleach minimally^28^. Under our illumination conditions, light pulses of 200 ms were sufficient to fully unblock the current (Fig. 3B), while shorter illumination times could be used to finely tune the degree of blockade. Once in the *cis* configuration, MAGu relaxes back to its thermodynamically stable *trans* state slowly, with a half-life of about 20 min in solution (Fig. S1B). Accordingly, relief of blockade persisted for many seconds in darkness after a brief flash of 380 nm light (Fig. 3C), eliminating the need for constant illumination, an important feature for future neurophysiology experiments.

**Figure 3.**
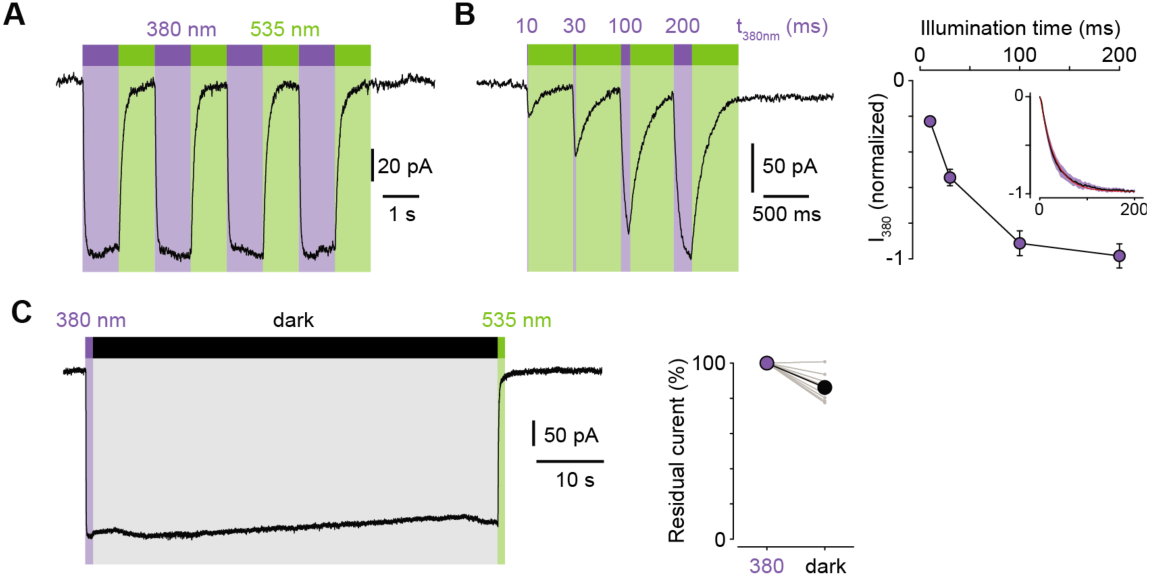
Photoregulation of GluD2-A654T-I677C labeled with MAGu. A. Representative recording showing the reversibility of block/unblock over multiple cycles of 380/535 nm light. B. Left, representative recording showing the extent of current unblock when varying the illumination time under violet light. Right, quantification of current unblock as a function of illumination time (n = 8 cells). Inset, averaged time course of current unblock when switching from dark to 380 nm light (mean value in black, ±SEM in purple, n = 7 cells) and corresponding mono-exponential fit (red, k = 0.0296 ± 0.0002 ms-1; t1/2 = 23.4 ms). C. Left, representative current trace showing the thermal stability of *cis* MAGu in darkness after a brief flash of 380 nm light. Right, 86.2 ± 2.5 % of the residual current remains after 1 min in darkness (n = 9 cells). Data are presented as mean value ± sem.

From a pharmacological point of view, current blockade occurred in the *trans* state (535 nm) and was relieved in the *cis* configuration (380 nm) for all membrane potential tested, with virtually no voltage-dependence (Fig. 4A), which contrasts with the profound voltage-dependence of block observed with pentamidine. This suggested to us that the positive charge of MAGu may not sense the electrical field of the membrane as much as pentamidine does. To investigate whether MAGu and pentamidine compete for the same binding site, we evaluated the dose-response relationship of pentamidine block on GluD2-L654T-I677C conjugated with MAGu, under both 380 and 535 nm light (Fig. 4B). We found the IC50s under both wavelengths to be virtually indistinguishable, favoring the idea that MAGu and pentamidine have distinct, non-overlapping binding sites. To get further molecular insight into *trans* MAGu-induced reduction of current, we performed molecular modeling experiments. After inserting the cysteine mutation, *trans* MAGu was docked by covalent docking (see methods). We found that the guanidinium headgroup of *trans* MAGu couldn’t reach the lumen of GluD2 (Fig. 4C), in agreement with our electrophysiology data. The effect of *trans* MAGu on the ion current was tested with the MOLEonline webserver^29^. We found that the photoswitch has a direct steric effect on the size of the cavity above the channel, as shown by the comparison of the computed channel in presence or absence of the photoswitch (Fig. 4D). In addition, the charge of the photoswitch could modify the electrostatic potential in the cavity and thereby affect ion transfer.

**Figure 4.**
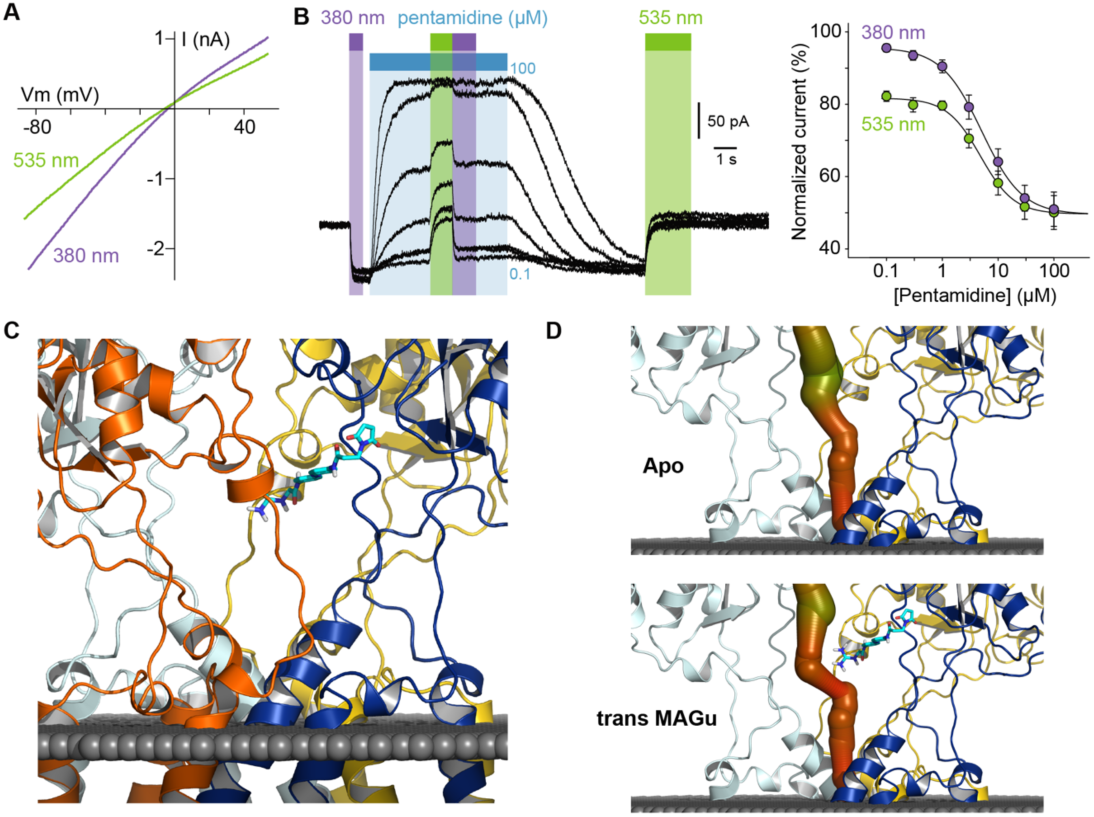
Pharmacological action of MAGu at GluD2-A654T-I677C. A. Representative current-versus-voltage relationship under 380 and 535 nm light. B. Left, representative dose-dependent blockade of the current upon pentamidine (0.1-100 μM) application, under 380 and 535 nm light. Right, quantification of pentamidine blockade under both wavelengths of light. IC50_380_ = 5.1 ± 0.3 μM, IC50_535_ = 4.8 ± 0.5 μM (n = 6 cells). Data are presented as mean value ± sem. C. Molecular modeling showing *trans* MAGu tethered to I677C. D. Molecular modeling showing the ion channel computed in the absence (top) and presence (bottom) of *trans* MAGu. The channel is represented with a color coding of the diameter in order to facilitate observation of the change induced by *trans* MAGu. One subunit is omitted for clarity.

We next sought to determine whether the non-Lc, native channel could be photocontrolled after installation of MAGu on the cysteine-substituted GluD2-I677C receptor. In heterologous expression system, activation of mGlu1 using the selective agonist 3,5-Dihydroxyphenylglycine (DHPG) was reported to trigger opening of GluD2 receptors ^20,21^. Therefore, we co-expressed the b isoform of mGlu1, which displays low basal activity^30^, together with GluD2 in HEK cells. Cells were labeled with MAGu and DHPG currents were recorded while alternating between 380 and 535 nm light. We found that DHPG-induced currents were reversibly reduced by about 23 % under 535 nm compared to 380 nm light for I677C, indicating that optical blockade with MAGu could be transposed to the native, non-Lc GluD2 (Fig. 5A). Importantly, DHPG-induced currents were identical in both wavelengths of light for the WT receptor (Fig. 5B), confirming that the effect of light is specific to the attachment of MAGu to I677C (Fig. 5C). In addition, we observed that the holding current increased when switching from darkness to 380 nm light for I677C, and decreased when switching back to 535 nm light, but remained constant in both wavelengths of light for WT (Fig. 5D). This suggests that a fraction GluD2 receptors are open prior to DHPG application, likely due to some basal mGlu1 activity in these cells. Altogether, these results show that the GluD2 I677C mutant labeled with MAGu (a.k.a. LiGluD2) possesses a functional ion channel, which can be gated through the mGlu signaling pathway, and which can be reversibly blocked and unblocked with green and purple light, respectively.

**Figure 5.**
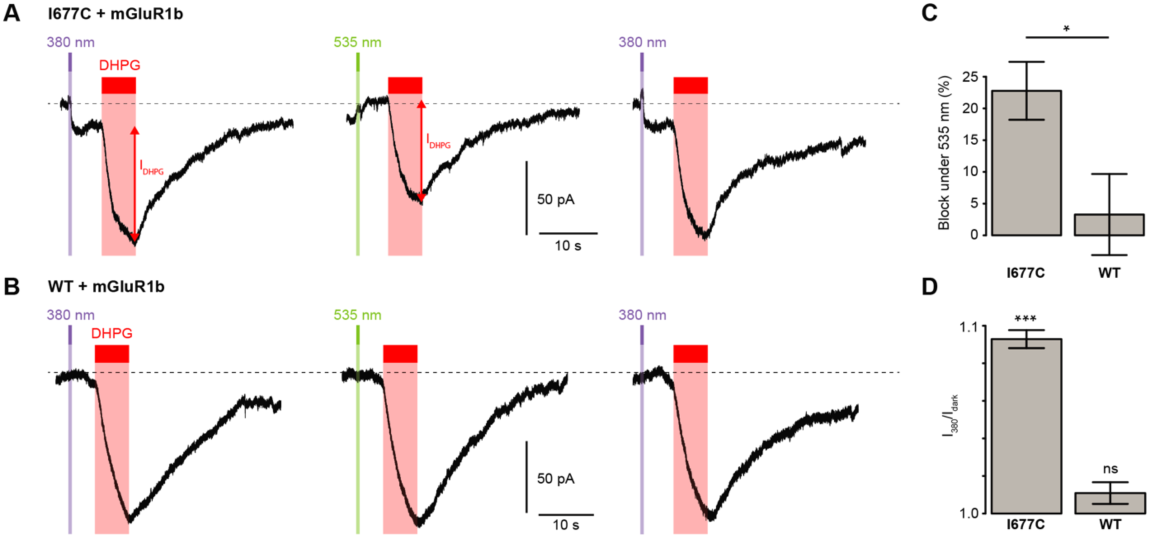
Photocontrol of GluD2-I677C (LiGluD2). A. Representative DHPG-induced current for a MAGu-treated (20 µM, 20min) cell co-expressing mGlu1b and GluD2-I677C, under 380, 535 and 380nm light. Note the drop in holding current at the onset of the 380-nm illumination, and the return to the baseline under 535 nm light. B. Representative DHPG-induced current for a MAGu-treated cell co-expressing mGlu1b and WT GluD2, under 380, 535 and 380nm light. C. DHPG-induced currents were reduced under 535 compared to 380 nm light for I677C (22.8 ± 4.6 %, n = 8 cells) but not for WT GluD2 (3.3 ± 6.4 %, n = 6 cells, p = 0.02, two-sample t-test). D. Ratio of the holding current recorded under 380 nm light and in darkness is different from 1 for I677C (1.09 ± 0.004 %, p = 2.44 e-7, n = 8 cells) but not for WT GluD2 (1.01 ± 0.006 %, p = 0.10, n = 8 cells, one-sample t-test).

The PTL strategy has been successfully applied to several members of the iGluR family, including kainate^31^ and NMDA^32^ receptors. In these former studies, the photoswitches were made with a glutamate head group, and were tethered to the LBD in proximity to the glutamate binding pocket, providing photocontrol of channel gating^24^. Because it remains unclear whether ligands can directly trigger GluD activation upon binding to the LBD, we adopted a different strategy for photocontrolling GluD receptors. We installed the photoswitchable ligand MAGu in proximity to the pore lumen, in hope to alter ion conduction through non-competitive antagonism. We found several cysteine mutants for which current was specifically modulated by light after attachment of MAGu, notably I677C (a.k.a. LiGluD2). In LiGluD2, *trans* MAGu likely does not reach the pore lumen as originally designed. Nevertheless, it reversibly and potently modulates current through the open GluD2 channel with high temporal and pharmacological precision.

The compounds traditionally used to probe the ionic function of GluDs, such as pentamidine and NASPM^16,17^, are not specific of GluD and also block NMDA and AMPA receptors. As to D-serine and glycine, they partially inhibit GluD2^Lc^ and mGlu1-gated GluD currents^13,20,22^, but they are also co-agonists of NMDA receptors. Here, the pharmacological specificity of LiGluD2 is absolute: the WT receptor is insensitive to either MAGu or light. Likewise, MAGu was shown to not photosensitize native brain tissue^25^. In fact, the PTL approach has already demonstrated exquisite pharmacological specificity for a large variety of cysteine-substituted ion channels and receptors^24,33^, even in complex biological settings such as brain slices^32^ or intact neuronal circuits *in vivo*^25,34^ where cells harbor free extracellular cysteines. Attachment of MAGu to GluD2 requires a single amino acid substitution, which is unlikely to disrupt the function of the receptor. In line with this, we found that the functional coupling of GluD2 with mGlu1 signaling^20,21^ was intact in LiGluD2. This enabled us to demonstrate with absolute pharmacological specificity that activation of mGlu1 triggers the opening of the GluD2 channel in heterologous expression system, in support of earlier evidence that the ion channel of GluD receptors may be functional^20-23^. LiGluD2 remains to be deployed in neuronal setting, yet we believe it will be a crucial tool for probing the ionotropic contribution of this orphan receptor to synaptic physiology.

## Materials and methods

### Chemicals

Bio-grade Chemicals products was provided by Sigma-Aldrich from Merck. MAGu was synthesized as previously described^25^ and provided by Enamine Ltd., Kyiv, Ukraine (www.enamine.net). MAGu was stored at −80°C as stock solutions in anhydrous DMSO.

### Spectrophotometry

UV-visible spectra were recorded on a Nanodrop 2000 (Thermo Scientific, 1 mm path) with 100 μM MAGu in PBS pH 7.4 (10% final DMSO). The sample was illuminated for 1 minute using ultra high-power LEDs (Prizmatix) connected to an optical fiber (URT, 1 mm core, Thorlabs), followed by an immediate measurement of absorbance. Light intensity at the tip of the 1 mm fiber was 100 mW for the 390 nm LED, and 150 mW for the 520 nm LED.

### Molecular biology

The single-cysteine mutations of GuD2 were generated by site-directed mutagenesis using the Quick Change II kit (Agilent technology) performed on pcDNA3-GluD2^20^. All mutants were verified by sequencing.

### Cell culture

Human Embryonic Kidney cells (HEK tsA201) were cultured in 25 cm^2^ tissue culture flask (Falcon, Vented Cap, 353109) with a culture medium composed of Dulbeco’s Modified Eagle Medium (Gibco life technologies, 31966047) containing Glutamax and supplemented with Fetal Bovine Serum (10%, Gibco life technologies, 10500064), Nonessential Amino-Acids (1%, Life Technologies, 11140–035), ampicillin, streptomycin (50,000 U, Gibco, life technologies, 15140-122) and mycoplasma prophylactic (2.5 mg, InvivoGen) antibiotics.

### Transfection

HEK tsA201 cells were freshly seeded and plated out in a 6-well plate, on coverslips (10 mm) treated with poly-L-lysine hydrobromide (Sigma, P6282-5MG). Cells were transiently transfected using calcium-phosphate precipitation, as described in^35^, using 1 µg of cDNA of GluD2 cysteine mutant per well. For co-transfection experiments, we used mGlu1b/GluD2 ratio from 0.7 to 1, with a maximum of 2 µg of total DNA. The plasmid pRK5-mGlu1b used in this study is a generous gift of L. Prezeau (IGF, Montpellier)

### Electrophysiology

Electrophysiological currents were recorded on HEK tsA201 cells at room temperature (21-25°C), 24-48h after transfection. Prior to whole cell patch-clamp experiments, cells were incubated for 20 minutes with an extracellular solution containing 20 μM MAGu, and then washed for at least 5 minutes with a fresh external solution. Cells were perfused with an external solution containing (in mM): 140 NaCl, 2.8 KCl, 2 CaCl2, 2 MgCl2, 12 glucose, 10 HEPES and NaOH-buffered at pH 7.32. Cells were patched with a borosilicate pipette (4-5 MΩ) containing an intracellular solution containing (in mM): 140 KCl, 5 MgCl2, 5 EGTA, 10 HEPES, and pH-adjusted to 7.32 with KOH. For recording metabotropic activation of GluD2 by mGlu1, the internal solution contained (in mM): 140 K-gluconate, 6 KCl, 12.6 NaCl, 0.1 CaCl2, 5 Mg-ATP, 0.4 Na-GTP, 1 EGTA, 10 HEPES, and was adjusted to pH 7.32 with KOH. Pentamidine solutions were applied using a fast-step perfusion system equipped with three square tubes (SF77B, warning instruments), as described in^35^. Illumination was carried out using a high-power LED system (pE-2, Cooled) mounted directly on the epifluorescence port of a vertical microscope (SliceScope Pro 6000, Scientifica). Currents were recorded with an axopatch 200B and digitized with a digidata 1440 (Molecular devices). Signals were low-pass filtered (Bessel, 2 kHz) and collected at 10 kHz using the data acquisition software pClamp 10.5 (Molecular Devices). Electrophysiological recordings were extracted using Clampfit (Molecular Devices) and analyzed with R.

### Molecular modeling

The model of the GluD receptor has been obtained by homology modeling using the software modeller version 9.19^36^. The template was that of the glutamate receptor GluA2 (PDB code 5weo)^27^. The automodel class has been used with slow level of MD refinement and the optimisation has been repeated 3 times for each model. 500 models were prepared and the best, as assessed by the DOPE score, was retained for further studies.

The structure of the protein and ligand were converted to pdbqt files with the software open babel 2.4.1. Covalent docking was then performed with the software smina^37^. The box of 25*25*25 angstrom was defined manually to encompass the mutated residue and extend to axis of symmetry. Covalent docking forced the maleimide to be in direct contact with the SG atom of the cysteine with which it is shown experimentally to form a covalent bond.

### Data analysis

Data are plotted as mean ±SEM. Total number (n) of cells in each group and statistics used are indicated in figure and/or figure legend. Comparisons between means were performed using parametric tests (two-sample t-test, Normality always verified, Shapiro-Wilk test of normality). Homogeneity of variances was tested preliminarily and the t-tests were Welch-corrected accordingly. For comparison with theoretical values of 0 or 1, we performed either one-sample t-tests when Normality was verified, or a non-parametric test (one-sample Wilcoxon tests) when Normality was not verified. ^#^p<0.1, *p<0.05, **p<0.01, ***p<0.001.

Time-course of current unblock and of thermal relaxation were fitted with the following mono-exponential function:

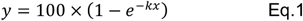

with k the time constant, and ln2/k the half-life.

Dose-response relationships were fitted with the following equation:

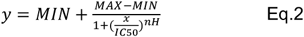

with MAX the maximal current, MIN the minimal current, IC50 the pentamidine concentration yielding half block, and nH the Hill number.

## Acknowledgements

Authors would like to thank Nadine Mouttajagane and Manel Badsi for their help with molecular biology work. This work was supported by funding provided by the French Agency for Research (ANR-16-CE16-0014-01 to LT, ANR-11-LABX-0011 to AT), by the “Initiative d’Excellence” (cluster of excellence LABEX Dynamo) to AT, by the Foundation for Medical Research (FRM, Equipe FRM EQU201903007961 to P.F) and by a post-doctoral fellowship from the Labex BioPsy to DL.

## Author contribution

Performed experiments: DL, SM and JT. Performed molecular modeling: AT. Performed analysis: AM and DL. Supervised the work: AM, LT, PF and BL. Wrote the paper: AM, BL and LT, with inputs from all the other authors. Acquired funding: PF, AT and LT.

