## Supplementary material for "Probing the ionotropic activity of the orphan glutamate delta 2 receptor with genetically-engineered photopharmacology": Supp. material

Figure S1

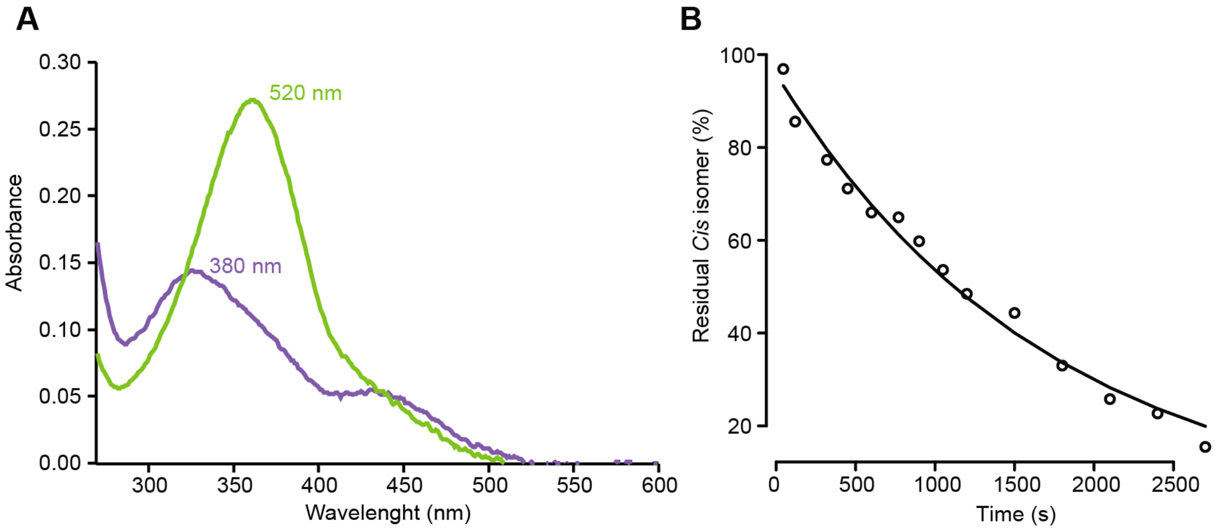


**Supplementary Figure 1.** Photochemical properties of MAGu. A. UV-visible spectra of MAGu under 520 nm light (green, mostly *trans*) or under 390 nm light (pink, mostly *cis*). B. Thermal relaxation of *cis* MAGu in aqueous solution in darkness. Absorbance at 362 nm is plotted as a function of time in darkness following illumination with 390 nm light. Data points were fitted with the following monoexponential decay equation: y = A exp(-x/k) and yielded: A = 95.8 ± 1.9 and k = 5.8e-04 ± 0.2e-04 $s^{-1}$. The half-life of *cis* MAGu was ln(2)/k = 1195 s.

### Figure S2

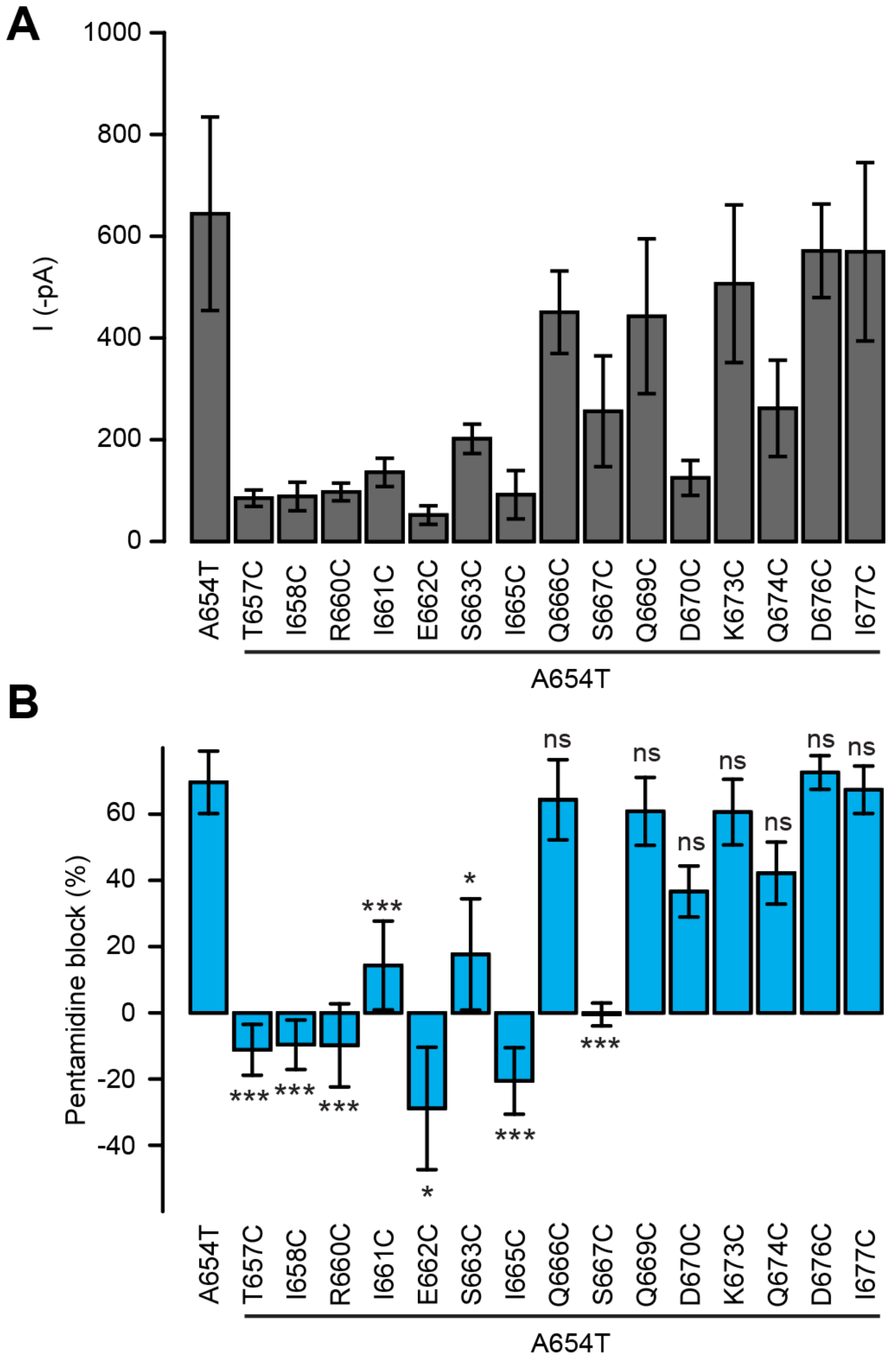


**Supplementary figure 2.** Functional characterization of the cysteine mutants. A) Currents recorded at -60 mV for A654T and each of the fifteen cysteine mutants (n = 3-8 cells). B. Pentamidine block for A654T and each of the fifteen cysteine mutants (n = 3-8 cells). Data are presented as mean value ± sem.
